# Oligodendrocyte Development and Myelin Sheath Formation are Regulated by the Antagonistic Interaction between the Rag-Ragulator Complex and TFEB

**DOI:** 10.1101/2023.03.02.530892

**Authors:** Ellen L. Bouchard, Ana M. Meireles, William S. Talbot

## Abstract

Myelination by oligodendrocytes is critical for fast axonal conduction and for the support and survival of neurons in the central nervous system. Recent studies have emphasized that myelination is plastic and that new myelin is formed throughout life. Nonetheless, the mechanisms that regulate the number, length, and location of myelin sheaths formed by individual oligodendrocytes are incompletely understood. Previous work showed that lysosomal transcription factor TFEB represses myelination by oligodendrocytes and that the RagA GTPase inhibits TFEB, but the step or steps of myelination in which TFEB plays a role have remained unclear. Here, we show that TFEB regulates oligodendrocyte differentiation and also controls the number and length of myelin sheaths formed by individual oligodendrocytes. In the dorsal spinal cord of *tfeb* mutants, individual oligodendrocytes produce fewer myelin sheaths, and these sheaths are longer than those produced by wildtype cells. Transmission electron microscopy shows that there are more myelinated axons in the dorsal spinal cord of *tfeb* mutants than in wildtype animals, but no significant change in axon diameter. In contrast to *tfeb* mutants, oligodendrocytes in *rraga* mutants produce shorter myelin sheaths. The sheath length in *rraga; tfeb* double mutants is not significantly different from wildtype, consistent with the antagonistic interaction between RagA and TFEB. Finally, we find that the GTPase activating protein Flcn and the RagCa and RagCb GTPases are also necessary for myelination by oligodendrocytes. These findings demonstrate that TFEB coordinates myelin sheath length and number during myelin formation in the central nervous system.

**Graphical Abstract:** 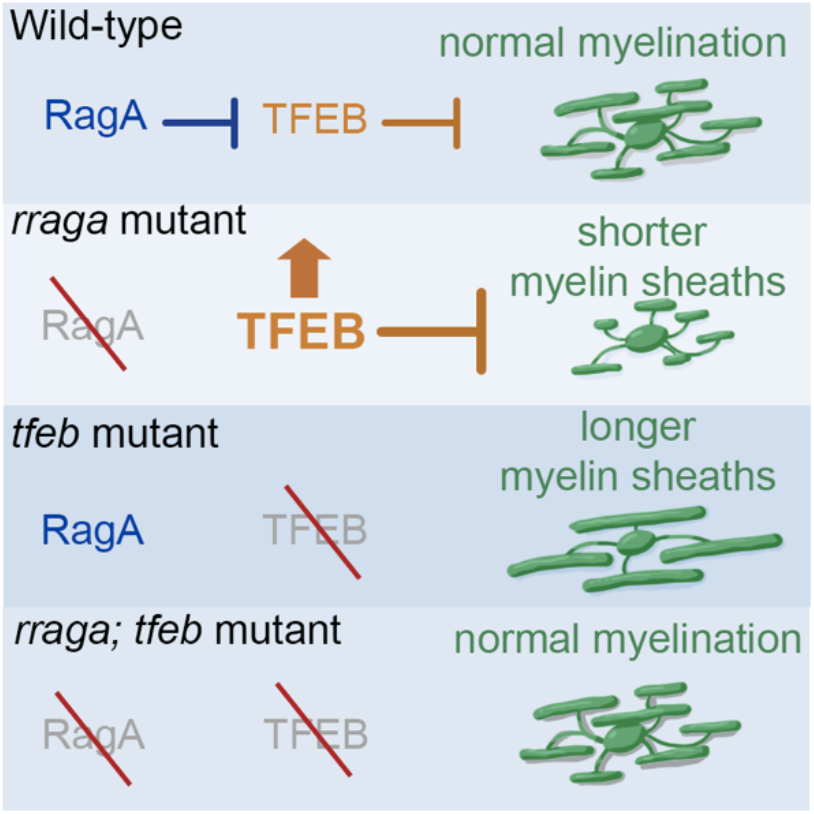

## Introduction

Myelin, the lipid-rich membranous sheath that surrounds axons, is critical for fast conduction of electrical signals in the nervous system and for the survival of myelinated axons (Fünfschilling et al., 2012; Sherman and Brophy, 2005; Simons and Nave, 2016). In the central nervous system (CNS), myelin is generated by oligodendrocytes (OLs). These myelinating glia differentiate from oligodendrocyte precursor cells (OPCs) during the development of the CNS. Some OPCs persist into adulthood and continue to give rise to myelinating OLs, contributing to new myelination in the adult CNS and to remyelination in disease (Emery, 2010; Lubetzki et al., 2020). New and changing myelination in the adult CNS is implicated in motor learning and memory (Bonetto et al., 2021; Chan and Xin, 2020; Foster et al., 2019). The importance of myelin is underscored by the demyelinating disease Multiple Sclerosis (MS), an autoimmune condition that disrupts CNS myelin and leads to impaired motor function, vision, and cognition (Browne et al., 2014; Dutta and Trapp, 2011; Franklin and Ffrench-Constant, 2008; Münzel and Williams, 2013). Disruptions to myelin have also been associated with pathology in models of Alzheimer’s disease (Chen et al., 2021; Maydell et al., 2022). Despite the critical functions of myelin in the central nervous system, our understanding of the molecular mechanisms that regulate myelination remains incomplete.

TFEB is well known as a regulator of lysosomal activity and autophagy (Napolitano and Ballabio, 2016; Sardiello, 2016; Sardiello et al., 2009). The guanosine triphosphate (GTP)-bound heterodimeric Rag-GTPases (RagA or B bound to RagC or D) integrate metabolic signals at the lysosome and regulate TFEB activity. In the presence of amino acids, RagA recruits TFEB to the lysosomal membrane, where it is phosphorylated and inactivated. In a starvation state, TFEB is dephosphorylated and translocates to the nucleus, where it activates the expression of genes related to lysosome biogenesis, autophagy, and lipid catabolism in order to maintain nutrient homeostasis (Martina and Puertollano, 2013; Settembre et al., 2011).

We previously reported that *rraga*, the gene that encodes RagA, is necessary for CNS myelination in zebrafish (Meireles et al., 2018). Simultaneous mutation of *tfeb*, the gene that encodes the transcription factor TFEB, rescues the hypomyelination phenotype in *rraga; tfeb* mutants. Moreover, transmission electron microscopy revealed more myelinated axons in the spinal cord of *tfeb* mutant animals than in control animals, indicating that TFEB represses myelination in the CNS. In rodents, TFEB controls the survival of pre-myelinating oligodendrocytes and also has an independent repressive effect on myelination (Sun et al., 2018). It remains unclear how TFEB represses myelination.

We have investigated the steps of oligodendrocyte differentiation and myelination that are disrupted in *rraga* and *tfeb* mutant zebrafish. We find that *rraga* mutation prevents the differentiation of oligodendrocyte precursor cells (OPCs) into OLs, and that the OLs that do differentiate in *rraga* mutants form abnormally short myelin sheaths. Mutation of *tfeb* rescues both of these defects in *rraga; tfeb* double mutants, and *tfeb* single mutants form abnormally long myelin sheaths. We confirm that *tfeb* mutation increases the number of myelinated axons in the dorsal spinal cord, and we find that the cross-sectional area of these myelinated axons remains unchanged. Finally, we report that Flcn and the Rag-GTPases RagCa and RagCb are essential for proper CNS myelination. Our findings reveal that myelin sheath length and oligodendrocyte differentiation are regulated by the antagonistic relationship between RagA and TFEB in the developing CNS.

## Methods

### Zebrafish Lines and Husbandry

Zebrafish embryos, larvae and adults were produced, grown, and maintained according to standard protocols approved by the Stanford University Institutional Animal Care and Use Committee. To obtain embryos and larvae used in experiments, adults 3-18 months of age were crossed. Adult density was maintained at 5-10 fish/L, with a 12hr light/12hr dark cycle and fish were fed twice daily. Water temperature was maintained at 28°C. Embryos and larvae were treated with 0.003% 1-phenyl-2-thiourea (PTU) to inhibit pigmentation, and they were anesthetized with 0.016% (w/v) Tricaine prior to experimental procedures. Published strains used in this study include: wildtype TL, *Tg(olig2:DsRed)* (Kucenas et al., 2008; Shin et al., 2003), *Tg(cldnk:GFP-CAAX*) (Münzel et al., 2012), *rraga*^st77^ (Shen et al., 2016), and *tfeb*^st120^ (Meireles et al., 2018). Genotyping was performed using standard conditions for PCR and restriction enzyme digestion. For *rraga* genotyping, primers used were 5’- TGGTTCAGGAGGATCAGAGAG-3’ and 5’-CAAAAGCAGACAATGCAAAAC-3’. PCR products were digested with the restriction enzyme MspA1I, which cleaves the WT allele. For *tfeb* genotyping, the primers used were 5’-GCTCATGCGGGACCAAATGC-3’ and 5’-GGTCACACTAACAAATGTGG-3’. PCR products were digested with Cac8I (R0579, New England Biolabs), which cuts the wildtype allele. The diagnostic fragments were distinguished by running the digested PCR product on a 3% agarose gel.

### CRISPR/Cas9 Genome Editing

sgRNAs were designed using CHOPCHOP (https://chopchop.rc.fas.harvard.edu/) (Labun et al., 2016; Montague et al., 2014), transcribed with T7 polymerase (E2040S, New England Biolabs) and purified using mirVana miRNA isolation kit (AM1560, Ambion). sgRNA target sites were: *rragca* (5’- GGTTTACTGTCGGAGGGAGA-3’); *rragcb* (5’-GGCTTCATCTAACGGTGTCG-3’); and *rragd* (5’- GGAAACCGCGGATCCTGCTCA-3’). Cas9 protein (Macrolab, Berkeley, http://qb3.berkeley.edu/macrolab/cas9-nls-purified-protein/) was injected together with 300 ng sgRNA into 1-cell stage embryos, and lesions were detected in genomic DNA of injected embryos by PCR and Sanger sequencing. Injected animals were raised to adulthood and outcrossed, and their offspring were genotyped for frameshift mutations. Fish carrying the following frameshift mutations were isolated: *rragca*^st164^ (+1bp), *rragcb*^st165^ (+7bp), and *rragd*^st166^ (+4bp). For genotyping, standard PCR amplification was performed with the following primers: *rragca:* 5’-GTCAACATGTCGATCCAGTACG-3’ and 5’- ATAGCTTGTTTGCAGCGACTC-3’; *rragcb:* 5’-GTGGTATAACAAATGTTGATACTGCC-3’ and 5’- CTTTTCCAGACTGGCGTCAGC-3’; *rragd:* 5’-GCCTTTATATAGTGCTGGGCTTC-3’ and 5’- GGGTCGAAGAAATCAATCTGAC-3’. Animals were genotyped by Sanger sequencing of PCR products from genomic DNA.

### Confocal Fluorescent Imaging and Analysis

Zebrafish embryos were mounted in 1.5% low melting point agarose in distilled water. Images were captured of a four-somite region of the spinal cord using a Zeiss LSM confocal microscope, with a Plan-Neofluar 10× objective (numerical aperture 0.30). CZI image files were analyzed using FIJI image analysis software. For OPC number, dsRed+ cells residing in the dorsal spinal cord were counted. For OL number, GFP+ cells residing in the dorsal spinal cord were counted. For myelin sheath analysis, the length of each myelin sheath associated with a GFP+ dorsal cell was measured. Ventral OL’s were excluded from the analysis as they develop too densely to be able to perform measurements on individual cells and myelin sheaths. Data from individual cells were pooled by genotype and analyzed in RStudio to quantify the average sheath length per cell, average number of sheaths per cell, total length of myelin per cell, and length of each individual sheath (Lysko and Talbot, 2022). Data were collected in three to four separate experiments and pooled.

### *In situ* Hybridization

*In situ* hybridization on embryos and larvae was performed using standard methods (Thisse et al., 2004). Briefly, embryos were fixed overnight in 4% paraformaldehyde, dehydrated for at least 2 hr in 100% methanol, rehydrated in PBS, permeabilized with proteinase K, and incubated overnight with antisense riboprobes at 65°C. The probe was detected with an anti-digoxigenin antibody conjugated to alkaline phosphatase (11093274910, Sigma-Aldrich). Images were captured using the Zeiss AxioCam HRc camera with the AxioVision software. Antisense probes against *mbp* were previously described (Lyons et al., 2009).

### Transmission Electron Microscopy

TEM was performed as described previously (Lyons et al., 2008). Briefly, decapitated embryo torsos were fixed in 2% glutaraldehyde and 4% paraformaldehyde in 0.1 M sodium cacodylate buffer (pH 7.4). The anterior portion of the larvae were used to isolate DNA for genotyping. For secondary fixation, samples were fixed in 2% osmium tetraoxide, 0.1 M imidazole in 0.1 M sodium cacodylate (pH7.4), stained with saturated uranyl acetate, and dehydrated in ethanol and acetone. Fixation and dehydration were accelerated using the PELCO 3470 Multirange Laboratory Microwave System (Pelco) at 15°C. Samples were then incubated in 50% Epon/50% acetone overnight, followed by 100% Epon for 4 hr at room temperature. Samples were then embedded in 100% Epon and baked for 48 hr at 60°C. Blocks were sectioned using a Leica Ultramicrotome. Thick sections (500–1,000 micrometers) for toluidine blue staining were collected on glass slides, stained at 60°C for 5s, and imaged with the LeicaDM2000 microscope using the Leica DFC290 HD camera and Leica Application Suite software. After the desired region of the spinal cord was reached, ultrathin sections were collected for TEM analysis on copper grids, were stained with uranyl acetate and Sato’s lead stain (1% lead citrate, 1% lead acetate, and 1% lead nitrate). Sections were imaged on a JEOL JEM-1400 transmission electron microscope.

### Quantification and Statistical Analysis

Sample sizes are indicated in each figure and/or figure legend. Zebrafish larvae were genotyped after image acquisition and data collection. In some experiments, heterozygous animals were excluded from the analysis. The experiments were not randomized. Sample sizes, mean ± standard deviation, statistical test, and P values are indicated in the figures or figure legends. Statistical significance was assigned at P < 0.05. Statistical tests were performed using GraphPad Prism 9 software.

## Results

### Differentiation of OPCs requires inhibition of TFEB by RagA

To analyze the role of RagA in OPC and OL development, we employed two transgenes that label cells of the oligodendrocyte lineage at different stages. The *olig2:DsRed* transgene labels neuronal progenitors and OPCs, whereas the *claudinK:GFP-CAAX* transgene encodes a membrane-bound GFP that labels mature OLs and their myelin sheaths (Münzel et al., 2012; Shin et al., 2003). To test the possibility that RagA is necessary for the proliferation or migration of OPCs in the pre-myelinated CNS, we examined *rraga* mutant animals expressing *olig2:DsRed*. OPCs were distinguished based on their location in the dorsal spinal cord (further dorsal than the normal ventral position of neural progenitors) in the region of somites 5-13. At 4 dpf, when myelination is underway in the zebrafish CNS (Preston and Macklin, 2015), *rraga* mutant animals had fewer dorsal *olig2:DsRed*+ cells in the spinal cord than their wildtype siblings (Figure 1B,D).

**Figure 1.**
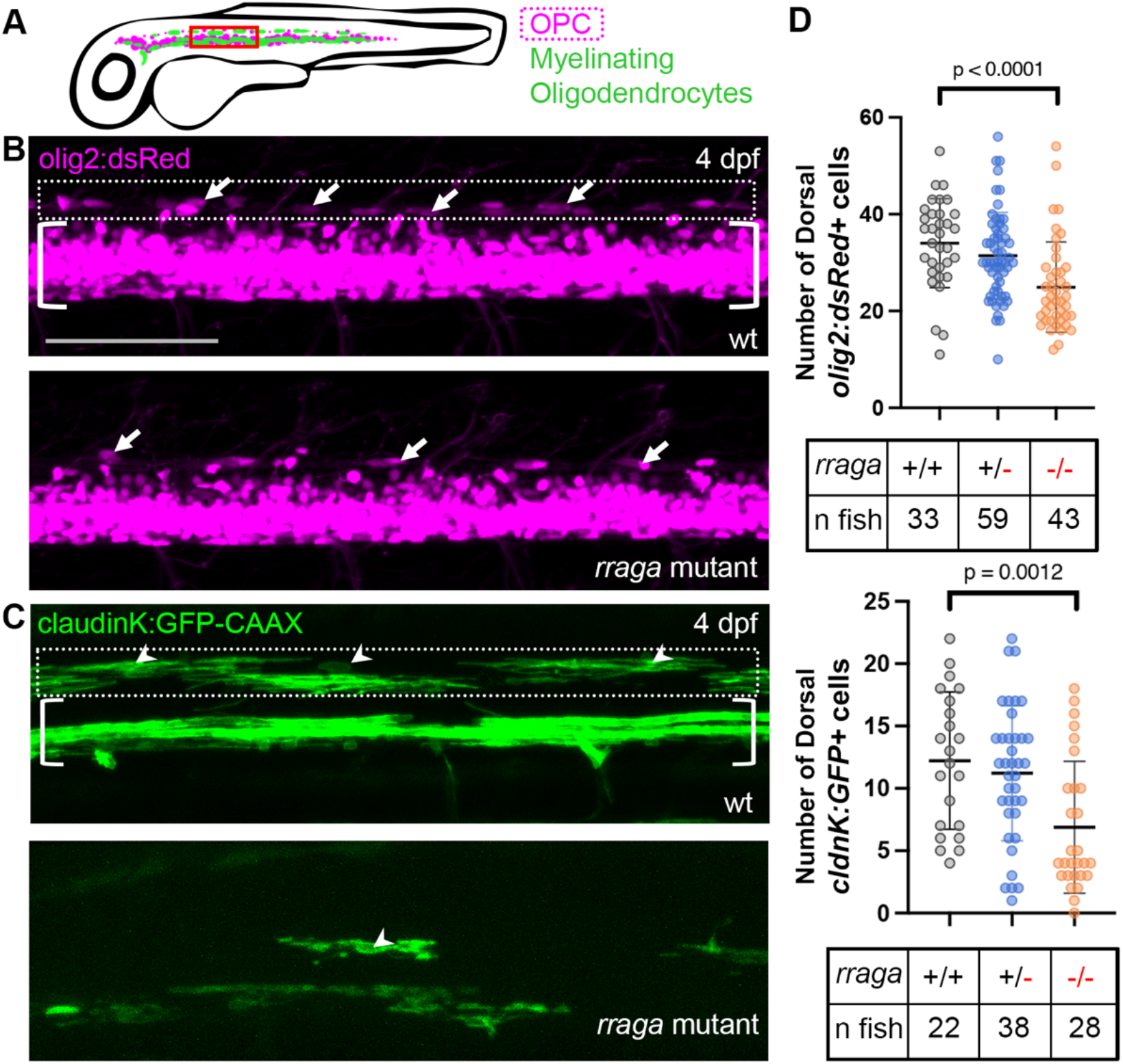
*rraga* is required for normal numbers of OPCs and oligodendrocytes in the dorsal spinal cord. **(A)** Diagram of a 4 dpf zebrafish larva, lateral view. The approximate patterns of olig2:DsRed (magenta) and claudinK:GFP-CAAX (green) expression are indicated. Red box indicates the 4-somite area shown in B & C. **(B)** Images of a 4-somite area of the spinal cord of olig2:DsRed-expressing animals at 4 dpf. Top: wildtype; bottom: *rraga* mutant. Dotted box indicates dorsal spinal cord region analyzed. Brackets indicate ventral spinal cord, which was excluded from analysis. **(C)** Images of a 4-somite area of the spinal cord of claudinK:GFP-CAAX-expressing animals at 4 dpf. Top: wildtype; bottom: *rraga* mutant. In B and C, arrows indicate dorsal OPCs. Dotted box indicates dorsal spinal cord region analyzed. Brackets indicate ventral spinal cord, which was excluded from analysis. Scale bar = 100μm. **(D)** Top: quantification dorsal OPC number in an 8-somite region of the spinal cord of wildtype (34.03±9.20), *rraga* heterozygous (31.41±8.97), and *rraga* mutant (24.93±9.33) animals at 4 dpf. Bottom: quantification of dorsal OL number in an 8-somite region of the spinal cord of wildtype (12.23±5.51), *rraga* heterozygous (11.21±5.40), and *rraga* mutant (6.89±5.29) animals at 4 dpf. Data aggregated from four separate experiments. P value indicates unpaired t-test with Welch’s correction. Comparisons that are not shown are not significant. Error bars indicate mean and standard deviation.

To determine whether the differentiation of OPCs into myelinating OLs is disrupted in *rraga* or *tfeb* mutants, we generated *rraga* mutant, *tfeb* mutant, and *rraga*; *tfeb* double mutant sibling animals expressing the membrane-bound OL marker *claudinK:GFP-CAAX*. We examined *claudinK:GFP-CAAX* expression in the dorsal spinal cord in the region of somites 5-13 at 4 dpf, where, on average, ∼35-40 *olig2:DsRed* expressing OPCs were present in wildtype embryos. *rraga* mutant embryos had many fewer *claudinK:GFP-CAAX+* OLs than wildtype controls (Figure 1C,D). These results show that the development and differentiation of OPCs into mature OLs is impaired in *rraga* mutant animals. By contrast, the number of OLs in *tfeb* single mutants is not significantly different from wildtype (Figure 2). In addition, *rraga*; *tfeb* double mutants have similar numbers of OLs as wildtype siblings (Figure 2), indicating that the loss of *tfeb* in *rraga; tfeb* double mutants rescues the disruption of OL differentiation in *rraga* mutants. Thus, our analysis indicates that RagA promotes OL differentiation by repressing TFEB.

**Figure 2.**
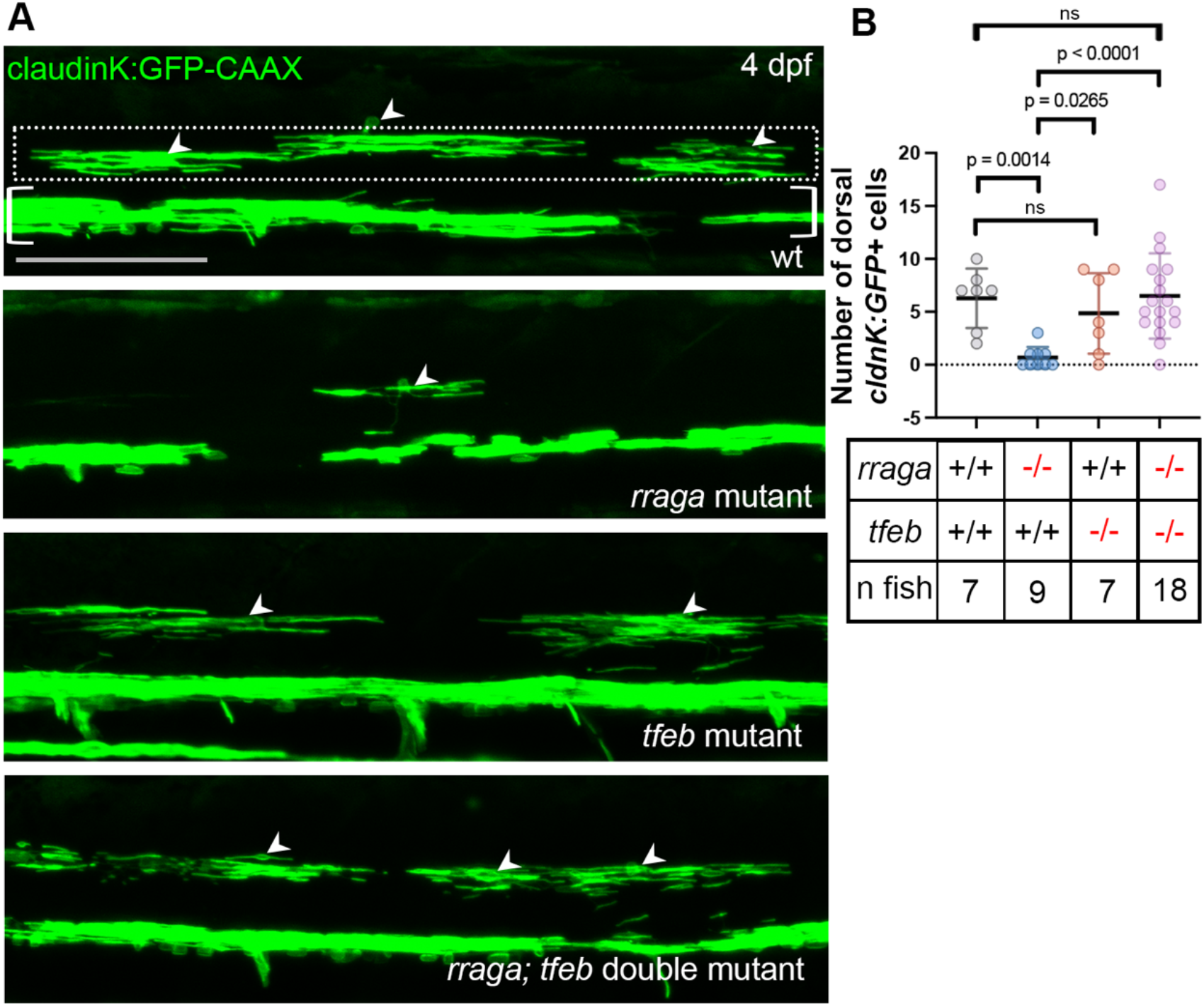
*rraga* promotes OL differentiation by repressing TFEB. **(A)** Images of 4-somite region of claudinK:GFP-CAAX-expressing animals at 4 dpf. From top down: wildtype, *rraga* mutant, *tfeb* mutant, *rraga; tfeb* double mutant. Arrowheads indicate dorsal myelinating oligodendrocytes. Dotted box indicates dorsal spinal cord region analyzed. Brackets indicate ventral spinal cord, which was excluded from analysis. Scale bar = 100μm. **(B)** Quantification of dorsal OL number in an 8-somite region of the spinal cord of wildtype (6.29±2.81), *rraga* mutant (0.67±1.00), *tfeb* mutant (4.86±3.81), and *rraga; tfeb* double mutant animals (6.50±4.03) at 4 dpf. Difference in number of OLs in *rraga* mutants between Figs. 2 and 3 reflect strain-to-strain phenotypic variation of *rraga* mutants. Data aggregated from three separate experiments. P value indicates unpaired t-test with Welch’s correction. Comparisons that are not shown are not significant. Error bars indicate mean and standard deviation.

### TFEB and RagA regulate myelin sheath length

Previously, we observed that *tfeb* mutant animals have more myelinated axons than wildtype siblings in the dorsal spinal cord at 9 dpf (Meireles et al., 2018), but it remains unknown how TFEB regulates myelin sheath formation. To determine how RagA and TFEB affect the parameters of myelin sheath formation by OLs, we measured myelin sheaths generated by individual OLs in the region of somites 5-13 in *rraga* mutant, *tfeb* mutant, and *rraga; tfeb* double mutant animals expressing *claudinK:GFP-CAAX* at 4 dpf (Figure 3A).

**Figure 3.**
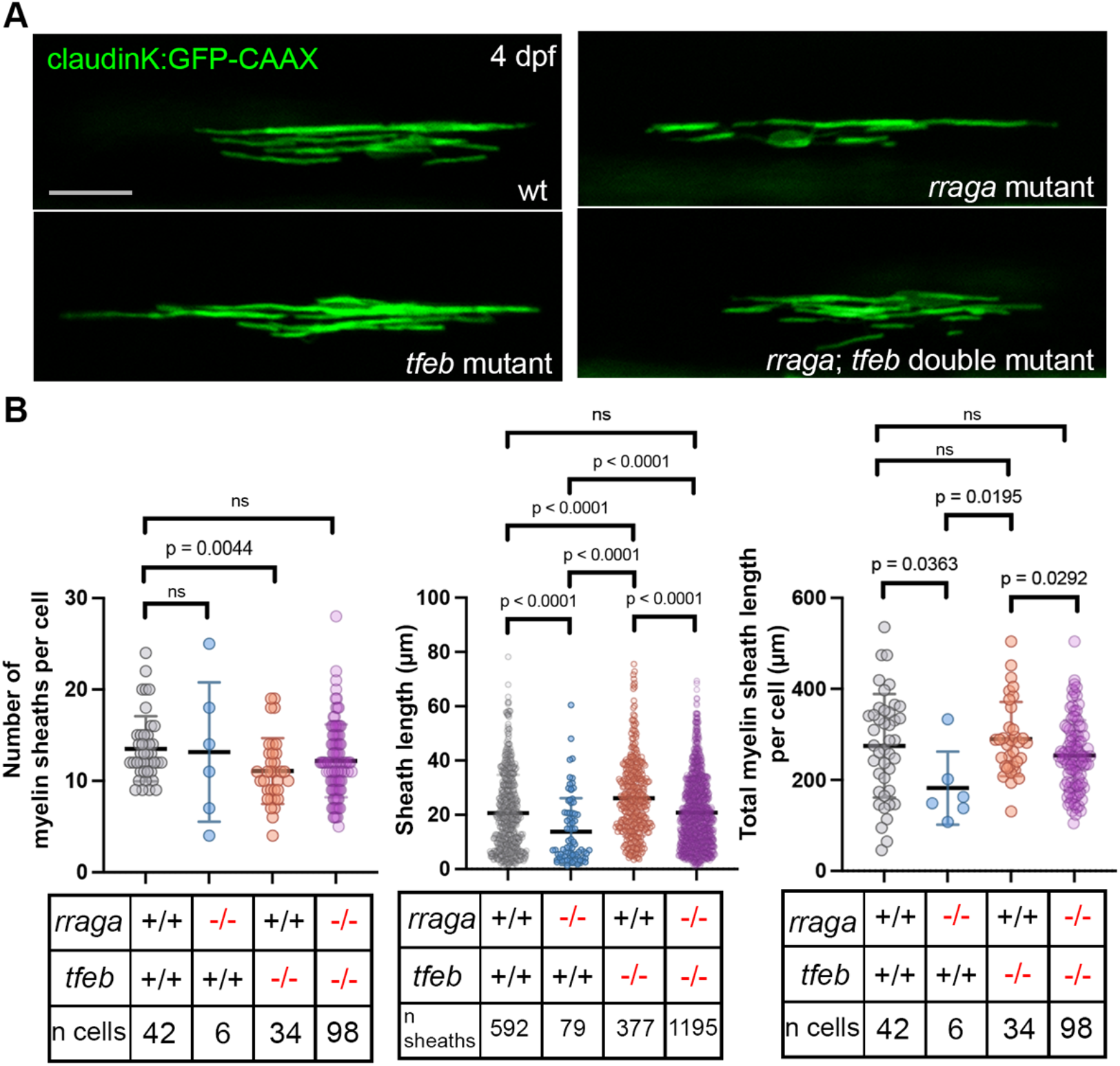
*rraga* and *tfeb* act antagonistically to regulate myelin sheath formation. **(A)** Images of single claudinK:GFP-CAAX-expressing oligodendrocytes in the dorsal spinal cord. Clockwise from top left: wildtype, *rraga* mutant, *rraga; tfeb* double mutant, and *tfeb* mutant. Scale bar = 25μm. **(B)** Left: quantification of number of myelin sheaths per cell in wildtype (13.52±3.56), *rraga* mutant (13.17±7.63), *tfeb* mutant (11.09±3.60), and *rraga; tfeb* double mutant (12.19±3.984) animals at 4 dpf. Middle: quantification of individual myelin sheath length in wildtype (20.61±14.09), *rraga* mutant (13.83±12.32), *tfeb* mutant (26.13±13.96), and *rraga; tfeb* double mutant (20.81±12.85) animals at 4 dpf. Right: quantification of total cumulative myelin sheath length per cell in wildtype (275.0±113.8), *rraga* mutant (182.0±80.26), *tfeb* mutant (289.7±81.59), and *rraga; tfeb* double mutant (253.8±77.28) animals at 4 dpf. P value indicates unpaired t-test with Welch’s correction. Data aggregated from three separate experiments. Comparisons that are not shown are not significant. Error bars indicate mean and standard deviation.

Although *rraga* mutant animals generate few myelinating OLs, a small number of *claudinK:GFP-CAAX*+ OLs were observed in the dorsal spinal cord and analyzed. The number of sheaths per OL was not significantly different in wildtype and *rraga* mutants, but the sheaths in *rraga* mutants were significantly shorter than in wildtype (Figure 3B). *rraga* mutant OLs also generated less total myelin, as measured by cumulative myelin length per cell, than wildtype controls (Figure 3B).

In *tfeb* single mutants, the number of sheaths per cell was slightly but significantly reduced, and the lengths of these sheaths were markedly increased compared to wildtype (Figure 3B). The total amount of myelin in *tfeb* mutants, as measured by cumulative length per cell, was not different from wildtype controls (Figure 3B). Finally, in *rraga; tfeb* double mutants, the number and length of sheaths was not significantly different than wildtype (Figure 3B).

These data indicate that myelin sheath number and length in the developing spinal cord are regulated by the antagonistic interaction between RagA and TFEB. Our observation that *tfeb* mutants have fewer, longer sheaths than wildtype indicates that TFEB influences OLs to produce shorter sheaths. RagA is essential to repress TFEB activity in myelinating OLs, such that *rraga* mutants have increased TFEB activity and reduced myelin sheath length.

### TFEB regulates myelination of dorsal axons independent of axonal cross-sectional area

There is a well-established correlation between axon diameter and myelination status, with larger-diameter axons being more likely to be myelinated (Sherman and Brophy, 2005). In order to determine whether changes in axon caliber may be related to the differences in myelination seen in *tfeb* mutants, we performed transmission electron microscopy on *rraga, tfeb*, and *rraga; tfeb* double mutant animals at 9 dpf. The number and cross-sectional area of myelinated axons in two dorsal quadrants of the spinal cord were measured in two animals per genotype (Figure 4A). Consistent with our previous findings (Meireles et al., 2018), *tfeb* mutant animals had more myelinated dorsal axons than wildtype or *rraga* mutant siblings, and the number of myelinated axons in *rraga; tfeb* mutants was not significantly different from wildtype (Figure 4B). Importantly, the cross-sectional area of myelinated axons in all mutants was similar to wildtype (Figure 4B), indicating that the increase in myelination is not simply an effect of increased axon diameter.

**Figure 4.**
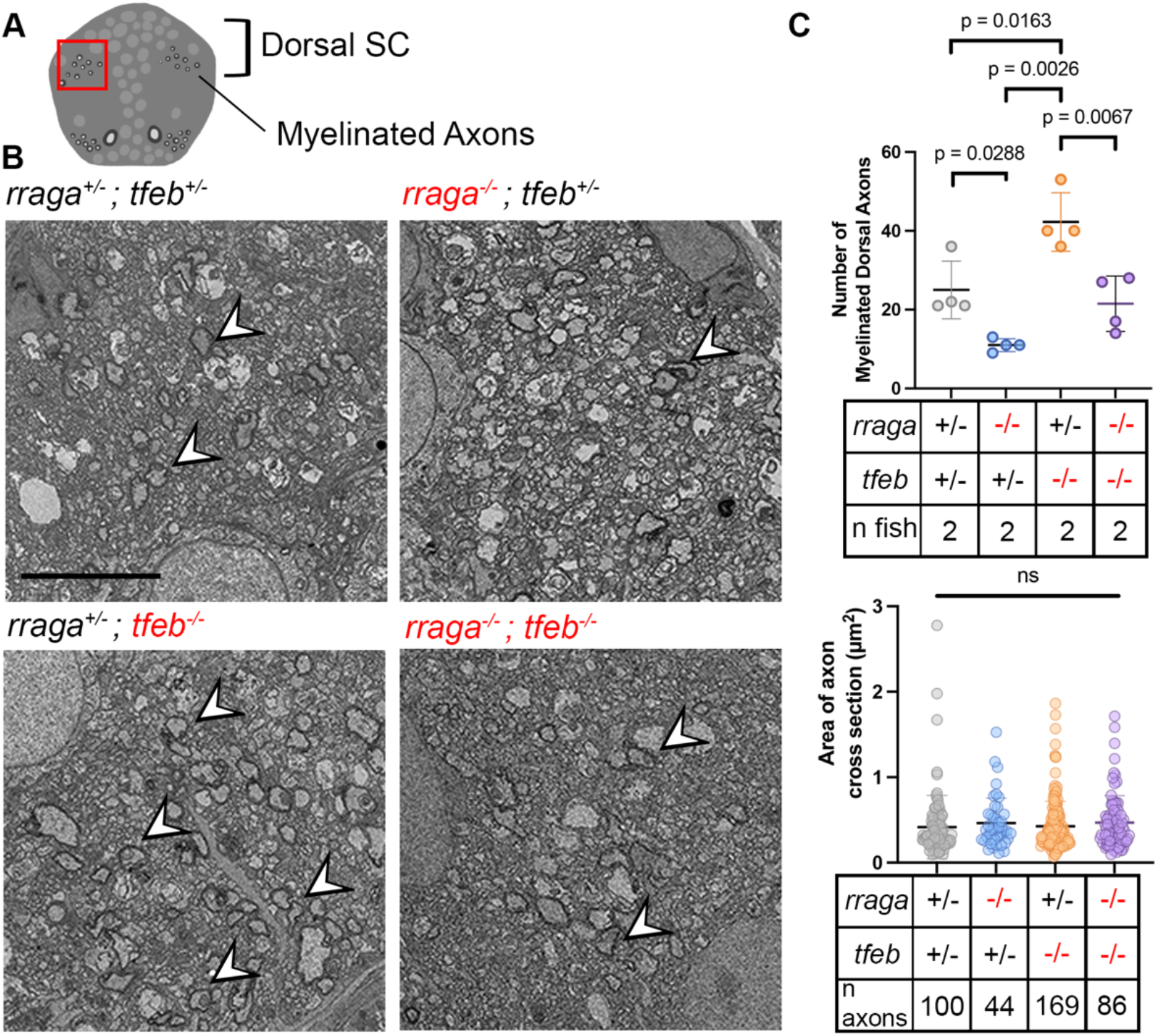
The number, but not cross-sectional area, of myelinated axons is increased in *tfeb* mutants. **(A)** Diagram of a cross-sectional view of the zebrafish spinal cord at 9 dpf. Bracket indicates area of dorsal spinal cord. Red box indicates dorsal quadrant area represented in B. **(B)** TEM images showing cross sections of the dorsal spinal cord in 9 dpf larvae. Clockwise from top left: control, *rraga* mutant, *rraga; tfeb* double mutant, *tfeb* mutant. White arrowheads indicate myelinated axons. Scale bar = 5μm. **(C)** Top: quantification of the number of myelinated axons per dorsal quadrant of the spinal cord in control (25.0±7.35), *rraga* mutant (11.0±1.63), *tfeb* mutant (42.25±7.41), and *rraga; tfeb* double mutant (21.5±7.05) animals at 9 dpf. Bottom: quantification of cross-sectional area of dorsal myelinated axons in control (0.416±0.370), *rraga* mutant (0.463±0.294), *tfeb* mutant (0.425±0.293), and *rraga; tfeb* double mutant (0.470±0.315) animals at 9 dpf. P value indicates unpaired t-test with Welch’s correction. Comparisons that are not shown are not significant. Error bars indicate mean and standard deviation.

### Lysosomal proteins RagCa, RagCb, and Flcn are essential for myelination by oligodendrocytes

In order to determine if other genes in the TFEB-related lysosomal network are essential for myelination, we investigated *flcn, rragca, rragcb*, and *rragd*. In mammals, the proteins RagA and RagB interchangeably heterodimerize with proteins RagC and RagD (Sekiguchi et al., 2001). Zebrafish lack an ortholog of the *rragb* gene, and there are duplicate genes encoding RagC, *rragca* and *rragcb*, such that RagA/Ca, RagA/Cb, and RagA/D are the putative functional heterodimers in zebrafish. The Flcn tumor suppressor has GAP activity for the RagC and RagD proteins, but not RagA or RagB (Tsun et al., 2013).

To examine the role of Flcn, we assessed *myelin basic protein* (*mbp*) mRNA expression by wholemount *in situ* hybridization in larvae from a cross of previously generated *flcn; tfeb* double heterozygotes (Iyer et al., 2022). At 5 dpf, *flcn* mutant animals had very little *mbp* mRNA in the CNS—a phenotype similar to *rraga* mutants (Figure 5A) (Meireles et al., 2018). In contrast, expression of *mbp* mRNA was similar in *flcn; tfeb* double mutants and controls (Figure 5A). These results show that, like RagA, Flcn is essential for CNS myelination and is required to repress TFEB activity.

**Figure 5.**
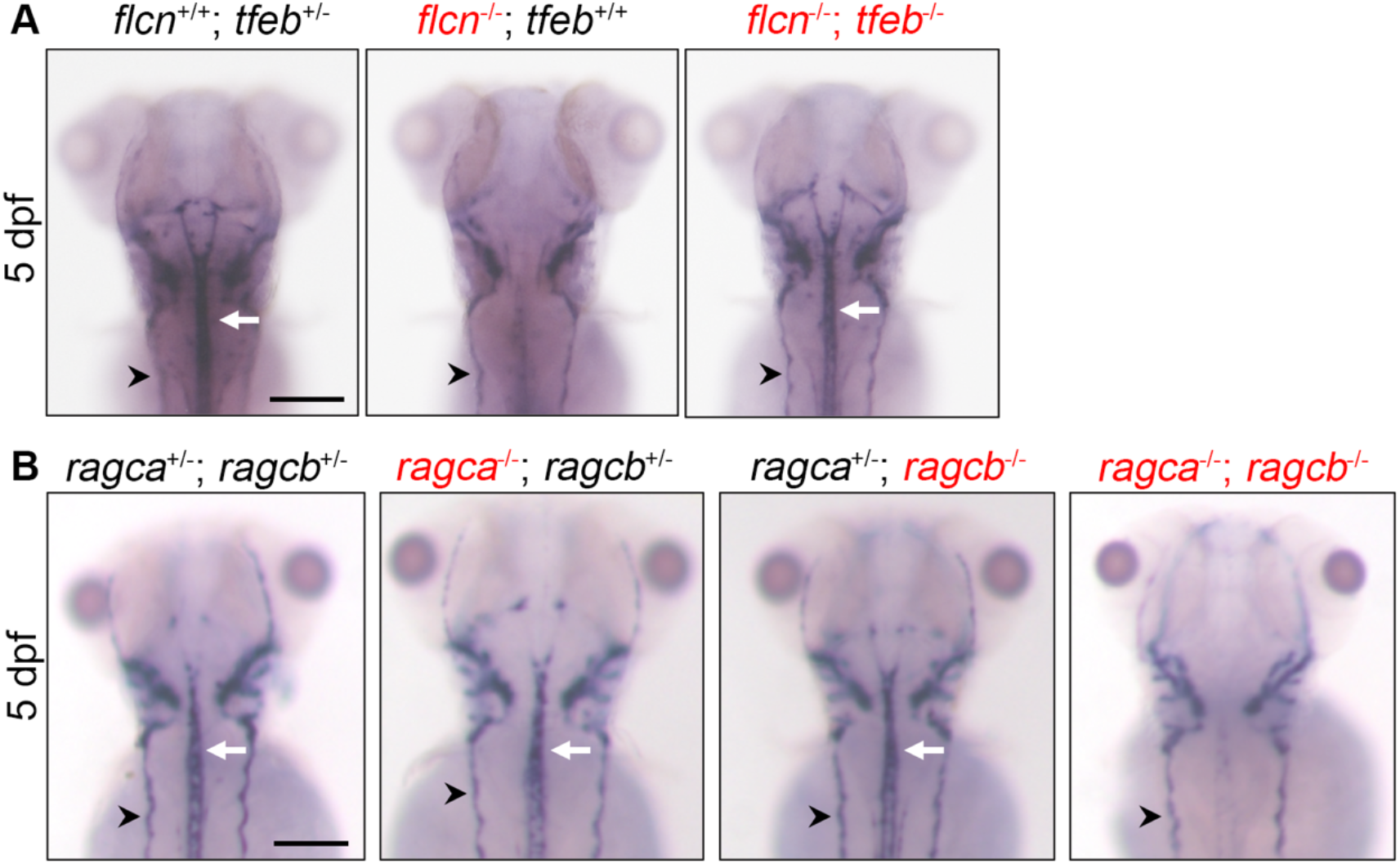
*flcn* and *rragca* and *rragcb* are essential for *mbp* expression in the CNS. **(A)** Images of 5 dpf zebrafish larvae stained for *mbp* mRNA via *in situ* hybridization. Left to right: control, *flcn* mutant, and *flcn;tfeb* double mutant. White arrows indicate CNS myelination in the spinal cord. Note reduced staining in the spinal cord of *flcn* mutant. Black arrowheads indicate PNS myelination. Scale bar = 50μm. **(B)** Images of 5 dpf zebrafish larvae stained for *mbp* mRNA via *in situ* hybridization. Left to right: control, *rragca* mutant, *rragcb* mutant, and *rragca;rragcb* double mutant. White arrows indicate CNS myelination in the spinal cord. Note reduced staining in the spinal cord of *rragca; rragcb* double mutant. Black arrowheads indicate PNS myelination.

Flcn activates the GTPase activity of RagC and RagD, so we also investigated the role of RagCa, RagCb, and RagD. We used CRISPR/Cas9 genome editing to generate fish lines that carry frameshift mutations in the *rragca, rracb*, and *rragd* genes (Supplemental Figure 1). We crossed double heterozygotes for *rragca* and *rragcb*, and we analyzed *mbp* mRNA by *in situ* hybridization in the resulting embryos at 5 dpf. Both *rragca* single mutants and *rragcb* single mutants had an *mbp* mRNA pattern that was similar to control siblings (Figure 5B). However, *rragca; rragcb* double mutants lacked *mbp* mRNA in the CNS (Figure 5B), a phenotype similar to *flcn* and *rraga* single mutants (Meireles et al., 2018). *rragd* mutation did not affect *mbp* mRNA expression in either a wildtype background or in a *rragca* or *rragcb* single mutant background (Supplemental Figure 2). These results indicate that RagCa and RagCb act redundantly, such that function of at least one is necessary for CNS myelination.

## Discussion

Together with our previous work (Meireles et al., 2018), our current study provides strong evidence that a RagA-RagC heterodimeric GTPase is essential for myelination in the zebrafish CNS. Our genetic analysis demonstrates that *rragca; rragcb* double mutants have pronounced reduction of *mbp* expression in the CNS, similar to the phenotype of *rraga* mutants. *mbp* expression is apparently normal in *rragca* and *rragcb* single mutants, indicating that these genes have redundant functions in myelination. In addition, our genetic studies indicate that the GTPase activating protein Flcn is also essential for normal *mbp* expression in the CNS, consistent with prior biochemical evidence that Flcn stimulates the GTPase activity of RagC (Tsun et al., 2013). The essential function of these proteins in myelination is to repress the activity of the transcription factor TFEB, as demonstrated by the restoration of *mbp* expression in *rraga; tfeb* double mutants and *flcn; tfeb* double mutants. A topic for future work will be to investigate the factors that regulate Flcn and Rag activity in developing oligodendrocytes.

Our results also demonstrate that the antagonism between RagA and TFEB has a previously unknown function in regulating the length of developing myelin sheaths in the spinal cord. Our measurements indicate that myelin sheaths in the CNS are abnormally short in *rraga* mutants, abnormally long in *tfeb* mutants, and normal in *rraga; tfeb* double mutants. The phenotype of *tfeb* single mutants indicates that TFEB activity represses myelin sheath length. Moreover, the phenotype of short myelin sheaths in *rraga* mutants indicates that excess TFEB activity can repress sheath length below normal wildtype levels, as our results suggest hyperactive TFEB is a central cause of defects in *rraga* mutants. Interestingly, *tfeb* single mutants have longer sheaths that are fewer in number than in wildtype, such that the total length of myelin per cell is similar in *tfeb* mutants and controls. Thus, elimination of TFEB activity increases sheath length but does not increase the total amount of myelin made by individual oligodendrocytes, perhaps reflecting nutritional or other constraints on the cell’s ability to produce additional myelin. It will be interesting to determine how TFEB interacts with other signaling pathways and factors, such as calcium signaling, that control myelin sheath length (Baraban et al., 2018; Krasnow et al., 2018; Swire et al., 2021). Simultaneous mutation of *tfeb* can rescue many phenotypes of *rraga* mutants, but there are some phenotypic differences between *tfeb* single mutants and *rraga; tfeb* double mutants. These differences point toward essential functions of RagA in addition to repressing TFEB activity. In microglia, for example, RagA and Flcn are essential to repress TFE3a and TFE3b (Iyer et al., 2022).

Our results also confirm and extend evidence that Rag GTPases and TFEB are essential regulators of OL differentiation. In *tfeb* mutants, we observed an increase in the number of myelinated axons in the dorsal spinal cord at 9 dpf, consistent with previous findings (Meireles et al., 2018). In addition, we found no difference in the diameter of myelinated dorsal axons in *tfeb* mutants, indicating that the changes in myelination are not simply a result of alterations in axonal diameter. Interestingly, although our analysis at an earlier stage (4 dpf) did not detect changes in the number of differentiating OLs or the total amount of myelin made by each, we observed that more axons are myelinated by electron microscopy at 9 dpf, suggesting that the phenotype of *tfeb* mutants becomes more pronounced as development progresses.

At early larval stages, *rraga* mutants have too few OPCs and an even stronger reduction in number of differentiated OLs. Oligodendrocyte number is normal in *rraga; tfeb* double mutants, adding to the evidence that disruptions in *rraga* mutant OLs are due to increased TFEB activity. It remains unclear from our analysis if the effect on OPC number is the result of changes in OPC proliferation, migration, or cell death. Sun *et al*. previously demonstrated that TFEB activity promotes apoptosis of pre-myelinating oligodendrocytes, and that *tfeb* mutation results in an excess number of myelinating OLs in the mouse brain (Sun et al., 2018). In our analysis of the zebrafish spinal cord, we did not see an increase in OL number in *tfeb* single mutants at 4 dpf. In mammals, a high percentage of oligodendrocyte lineage cells die as a normal part of CNS development (Barres et al., 1992; Trapp et al., 1997). In zebrafish, the role of cell death in development of the oligodendrocyte lineage has not been as thoroughly investigated, but there is little evidence of OPC cell death during the period of developmental myelination (Buckley et al., 2010; Keefe et al., 2020). Thus, the role of TFEB in controlling survival of developing oligodendrocytes may differ in rodents and fish.

A full understanding of the broader role of TFEB in myelination requires further study. Homozygous *tfeb* mutant zebrafish are viable to adulthood and reproduce normally, but CNS function in behavior has not been examined in these animals. It remains unclear how TFEB’s repressive effects on oligodendrocyte differentiation and sheath length integrate with other pathways that regulate myelination. That TFEB acts downstream of the Rag-Ragulator protein network suggests that TFEB may integrate metabolic status and stress responses, perhaps to coordinate how oligodendrocytes use their resources in myelin production. The effect of TFEB on myelination is evidently direct, because RagA and TFEB act autonomously in oligodendrocytes (Meireles et al., 2018), and the diameter of myelinated axons is unaffected. Further studies are needed to identify the target genes downstream of TFEB that mediate its effects on oligodendrocyte development and myelination.

## Acknowledgments

We thank Talbot laboratory members for helpful discussions and technical advice and Tuky K. Reyes and Chenelle Hill for fish maintenance. Transmission electron microscopy was performed with assistance from John J. Perrino in the Stanford University Cell Sciences Imaging Core Facility (RRID:SCR_017787). E.L.B. was supported by a fellowship from the National Science Foundation. W.S.T. is a Catherine R. Kennedy and Daniel L. Grossman Fellow in Human Biology. This work was supported by NIH grant R35NS111584.

