## Supplemental Figures for "Oligodendrocyte Development and Myelin Sheath Formation are Regulated by the Antagonistic Interaction between the Rag-Ragulator Complex and TFEB"

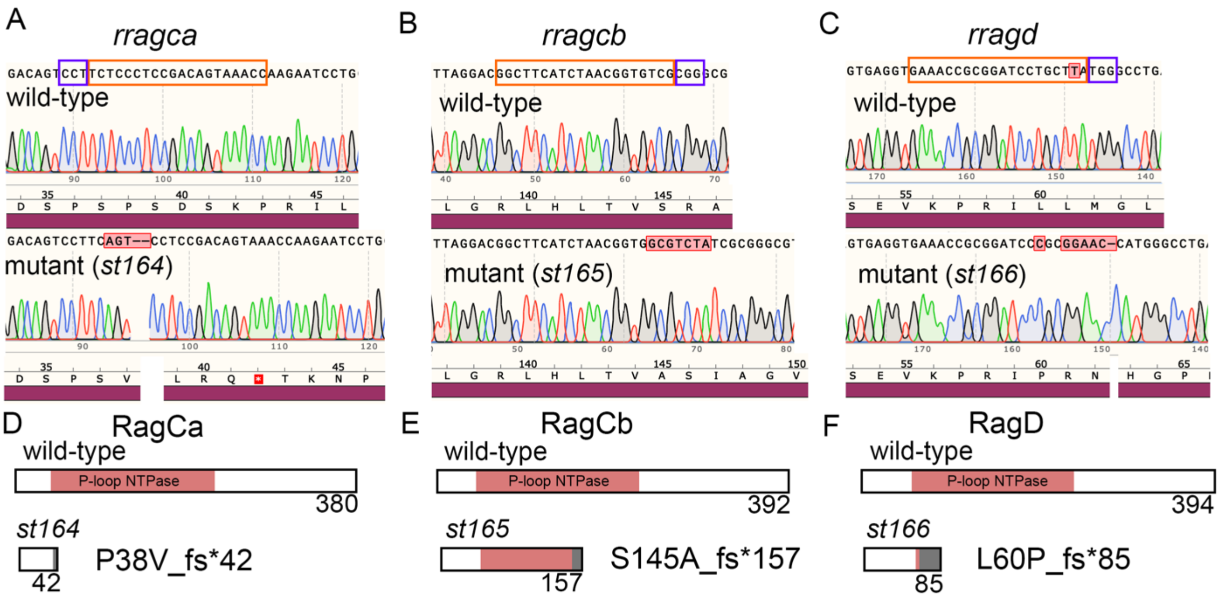


**Supplementary Figure 1. Effects of *rragca, rragcb, and rragd* mutations on predicted amino acid sequences.**

(A-C) Genomic DNA sequence at *rragca, rragcb, and rragd* lesion sites. Top: wildtype sequence and amino acid translation as aligned to reference sequence. Bottom: homozygous mutant sequence and amino acid translation as aligned to reference sequence. Orange box indicates sgRNA site, purple box indicates PAM site.

(D-F) Diagrams showing predicted effect of mutation on RagCa, RagCb, and RagD proteins. Top: wildtype protein. Bottom: predicted truncated protein product of mutant alleles. Red boxes indicate position of predicted P-loop containing NTPase domain. Grey boxes indicate disrupted amino acid sequence downstream of mutation site. Numbers indicate positions of stop codons. Bottom-right labels indicate amino acid change at mutation site as well as position of earliest stop codon. Diagrams are to scale.

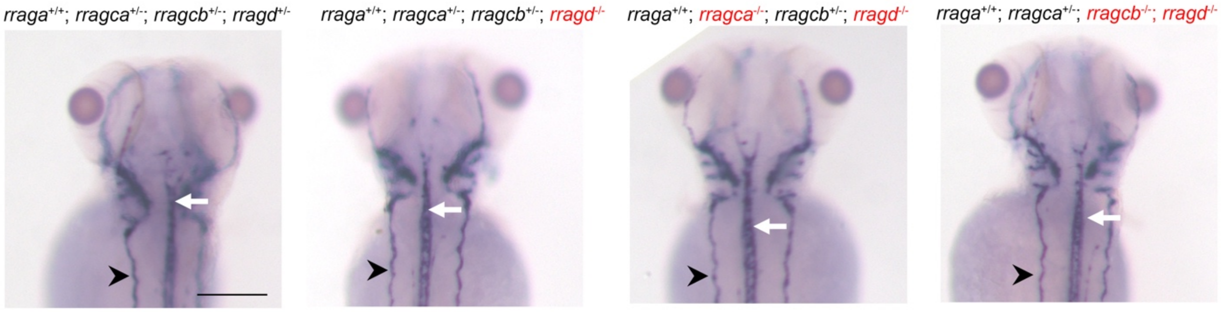


**Supplementary Figure 2. *rragd* mutation does not affect myelination in the CNS.**

Images of 5 dpf zebrafish larvae stained for *mbp* mRNA via *in situ* hybridization. Left to right: control, *rragd* mutant, *rragca; rragd* double mutant, and *rragcb; rragd* double mutant. White arrows indicate CNS myelination in the spinal cord. Black arrowheads indicate PNS myelination. Scale bar = 50µm.
